# Associations Between Shape Iconicity Ratings and Speech Acoustics: Comparing Real Words and Pseudowords

**DOI:** 10.64898/2026.05.19.725980

**Authors:** Josh Dorsi, Simon Lacey, Lynne C. Nygaard, K. Sathian

## Abstract

Iconicity in spoken language refers to the mapping of speech sounds to meaning. For example, the pseudoword “bouba” is judged to sound rounded, whereas the pseudoword “kiki” is judged to sound pointed (Ramachandran & Hubbard, 2001). Recent work has found systematic relationships between speech acoustics and iconicity ratings in various meaning domains, including shape. To control for confounding by semantic knowledge, much research on iconicity relies on pseudowords. However, the role of iconicity in conveying the meanings of real words is still poorly understood. Here, we compared the relationship between speech acoustics and shape iconicity ratings for both pseudowords and real words. In this experiment, participants rated real words and pseudowords for how rounded or pointed they sounded. We compared the relationship between 12 acoustic parameters and the rounded/pointed ratings for both words and pseudowords. We found that the correlations between these acoustic parameters and the ratings were similar for real words and pseudowords, providing key evidence that iconicity of real words is linked to speech acoustics.

---

Since Plato’s Cratylus dialogue (Ademollo, 2011), one of the oldest debates about language is whether the relationship between the sound of a word and its meaning is arbitrary (Weingartner, 1970). While the predominant view favors arbitrariness (Hockett, 1960; de Saussure, 1916/2009), many languages include words with both arbitrary and non-arbitrary sound-to-meaning relationships (Lockwood & Dingemanse, 2015; Blasi et al., 2016).

Iconicity in spoken language involves associations between speech sounds and sensory features or general concepts. It includes onomatopoeia, in which words mimic the sounds they refer to (e.g., "bark"; Catricalà & Guidi, 2015). Similarly, Japanese mimetic words resemble non-auditory stimuli; for example, "kirakira" refers to "flickering light" (Akita & Tsujimura, 2016). More generally, listeners report that certain words sound like their meaning (Winter et al., 2024) and that pseudowords are judged to sound rounded/pointed (Köhler, 1929, 1947), large/small (Sapir, 1929), rough/smooth (Greenberg & Jenkins, 1966), or dark/bright (Newman, 1933).

In the classic demonstration of speech iconicity, the pseudowords “maluma” and “takete” were ascribed to rounded and pointed shapes, respectively (Köhler, 1929, 1947; see also Ramachandran & Hubbard, 2001). Shape associations are the most widely studied instances of speech iconicity, with reports of shape correspondences for hundreds of different pseudowords (e.g., McCormick et al., 2015; Westbury et al., 2018). Much work has investigated the influence of phonetic structure on shape iconicity (D’Onofrio, 2014; Sidhu et al., 2021, 2022; Westbury et al., 2018). For example, pseudowords with voiced consonants tend to be associated with rounded shapes (Cuskley et al., 2017; D’Onofrio, 2014; Lacey et al., 2026; McCormick et al., 2015). Shape iconicity is also associated with vowel rounding (Maurer et al., 2006; McCormick et al., 2015; Lacey et al., 2026) and the presence of obstruents and sonorants (Ahlner & Zlatev, 2010; McCormick et al., 2015; Lacey et al., 2026).

Studying iconicity in pseudowords avoids the semantic confounds inherent in real words. For example, "balloon" *sounds* rounded (Sučević et al., 2015), but does knowing that balloons are round bias listeners’ judgments of the word’s sound? Partly for this reason, much of sound iconicity research relies on pseudowords. Implicit in this approach is the assumption that findings for pseudowords will generalize to real words. Indeed, there is some evidence that real words can carry iconic associations. For example, participant ratings of thousands of words indicate substantial counts of words that sound like their meaning (i.e., are iconic; see Winter et al., 2024, see also Monaghan et al., 2014). There also is notable consistency in certain sound-to-meaning mappings across many different languages (Blasi et al., 2016). Moreover, people are better than chance in correctly guessing the meanings of iconic words from unfamiliar languages (Lockwood et al., 2016). There is also evidence that iconic words from a foreign language are easier to learn than non-iconic words (Lockwood et al., 2016), and it is easier to learn the iconically congruent (relative to an arbitrary) meaning of a foreign word (Nygaard et al., 2009). By testing unfamiliar (i.e., foreign language) words, Lockwood et al. (2016) and Nygaard et al. (2009) isolated the sounds of words from their semantic associations. Foreign words are real words that are functionally pseudowords; thus, these learning effects suggest that iconicity is present in natural language.

There is also evidence that iconic associations for phonemes in pseudowords tend to occur in real words with matching meanings (e.g., Sidhu et al., 2021). Sidhu et al. (2021) asked participants to select the word with either the most rounded or most pointed meaning from sets of six orthographically presented words drawn from a corpus of over 1,700 words. Rounded/pointed *meaning* scores for each word were calculated from these participant responses. Drawing on prior work with pseudowords (Westbury et al., 2018), Sidhu et al. (2021) calculated a rounded/pointed *sound* score for each word based on its phoneme composition. Crucially, these two estimates converged: words with strongly rounded/pointed meanings also tended to have a high proportion of rounded/pointed phonemes, respectively (Sidhu et al., 2021).

There is also evidence linking speech acoustics to shape iconicity. For example, Parise and Pavani (2011), recorded participant vocalizations of a vowel while they were viewing different shapes: the third formant (F3)^1^ had a higher center frequency when speakers were viewing a triangle than when viewing a dodecahedron, possibly related to the triangle’s more prominently pointed features. In a perceptual task, Knoeferle et al. (2017) found that consonant-vowel pseudowords were rated as sounding more rounded than pointed if their vowel had a lower frequency F2 or (in contrast to the results of Parise & Pavani, 2011) a higher frequency F3. More recently, Lacey et al. (2020) found that rounded/pointed ratings for a large set of consonant-vowel-consonant-vowel (CVCV) auditory pseudowords (ratings collected by McCormick et al., 2015) correlated with several measures of vocal quality, such that greater acoustic variability was found to be associated with more pointed ratings. These relationships have been replicated (Kumar et al., 2025) and extended to a novel set of stimuli (Nayak, 2024), with additional evidence that shorter duration pseudowords tend to be rated as more pointed (Kumar et al., 2025). Lacey et al. (2020) also found associations between shape ratings and spectro-temporal properties of the speech signal — the fast Fourier transform (FFT), speech envelope, and spectral tilt — linking shape iconicity ratings to fundamental properties of the speech signal (replicated by Kumar et al., 2025 & Nayak, 2024).

A notable limitation of many iconicity investigations is the use of written stimuli (e.g., Monaghan et al., 2014; Blasi et al., 2016). Sidhu et al. (2021) collected participant judgments of word meaning that were based on orthographically presented words and compared those judgments to each word’s phonetic composition. While testing the relationship between meaning and phonology, this design begs the question of whether the findings can be generalized to the acoustics of verbal language. This is notable in light of recent work linking speech acoustics to iconic associations (e.g., Lacey et al. 2020; Kumar et al., 2025).

The present work builds on Lacey et al. (2020) by testing whether the acoustic-phonetic properties that support ratings of shape iconicity also characterize judgments of iconicity in real words. There are two reasons that these relationships might not generalize. First, they may be idiosyncratic to the pseudowords used in the prior studies whose stimuli were highly controlled to have a uniform syllabic structure (CVCV) and include a representative but not exhaustive sampling of English phonemes. This was ideal for observing the acoustic-to-rating relationships of interest to that work but raises the possibility that more word-like stimuli might not produce the same results. For example, while older children are able to match foreign language roundedness/pointedness words to rounded/pointed shapes they were more accurate when matching pseudowords designed to be iconic for roundness/pointedness to those shapes (Tzeng et al., 2017). Second, these relationships might depend on lexical status: they could characterize pseudowords but not real words. For instance, the correlations observed for pseudowords could arise because they lack semantic information on which to base shape judgments. This possibility is consistent with findings from outside the iconicity literature; for example, accuracy in speech in noise perception is better for real words than nonwords (Miller et al., 1951) and phoneme detection is faster for phonemes in real words than pseudowords (Rubin, Turvey, & Van Gelder, 1976) suggesting that lexical context can influence how phonemes are perceived.

We chose 160 real words from Sidhu et al. (2021) and constructed a set of 160 pseudowords derived from these real words to have comparable phonotactic probabilities. Additionally, a set of 320 CVCV pseudowords was drawn from the 537 pseudoword set used by Lacey et al. (2020) to provide a comparison to prior work. Because the derived pseudowords are more structurally similar to the real words than the CVCV pseudowords, these stimuli will be used as a strong test of the hypothesis that the ratings-to-acoustic relationship for pseudowords is consistent with that for real words. These comparisons allow us to account for the potentially more complex phonotactic structure of real words while still presenting items that lack semantic information.

## METHODS

### Participants

The data from 24 participants (20 female, 3 male, 1 non-binary; age: *M* = 30 years, *S* = 7) were analyzed for this experiment. A power analysis (IBM SPSS, Version 29) based on Kumar et al. (2025) data showed that this sample size afforded us more than 80% power to detect an acceptable degree of inter-rater reliability (Cronbach α of 0.8). Twenty-four participants also allowed a complete counterbalancing of ordering three stimulus blocks and two rating scales (see below). Data from one participant were discarded and replaced without being analyzed due to a computer error that prevented the participant from completing the experiment. All participants reported that American English was their first language and reported no hearing, speech, or language disorders. The Penn State University Institutional Review Board approved all procedures.

### Stimuli

This experiment used three types of speech stimuli: real words, pseudowords derived from these real words ("derived pseudowords"), and CVCV pseudowords. All stimuli were recorded in a random order by a female speaker whose first language was American English. Following the general procedures of McCormick et al. (2015), recordings were made in a sound-attenuated room using a Samsung C01U PRO microphone and were digitized at a 44.1 kHz sampling rate. Two independent judges assessed whether each item was recorded with a neutral intonation, sounded consistent with other recordings (e.g., was not louder/quieter, faster/slower), and contained the intended phonetic content. If either judge assessed that a given item did not meet these criteria, it was re-recorded and re-judged. Stimuli were amplitude-normalized using Audacity software (Mazzoni, 2006). Finally, stimuli were down-sampled to 22.05 kHz using Praat software (Boersma & Weenink, 2011).

### Real words

We used 160 real words (duration: *M* = 594 ms, *S* = 117 ms), a subset of the items presented as visual text by Sidhu et al. (2021). Sidhu et al. (2021) reported a measure of how rounded/pointed each word’s *meaning* was. We selected 40 words with very rounded and 40 with very pointed meaning scores. Words were selected through an iterative process: starting from the endpoints of the meaning-score distribution, we replaced items as needed so that the rounded and pointed sets were comparable on length and key lexical/phonotactic metrics. Next, we selected 80 words with meaning scores near either side of the median following a similar iterative process. This yielded two groups, one on either side of the median, corresponding to words relatively closer to the “rounded” and “pointed” ends. Thus, we had four groups of words: very rounded (n = 40), somewhat rounded (n = 40), somewhat pointed (n = 40), and very pointed (n = 40). These word groups did not differ in concreteness, lexical frequency, phonological neighborhood density, and biphone probability (see Supplemental Methods for definitions); however, they did differ in how rounded/pointed they would sound (using the "sound score" reported by Sidhu et al., 2021). Altogether, the 160 real words had between one and four syllables (median: 2; see Table 1). See the Supplemental Methods section for further details on the selection of the real words.

**Table 1.**
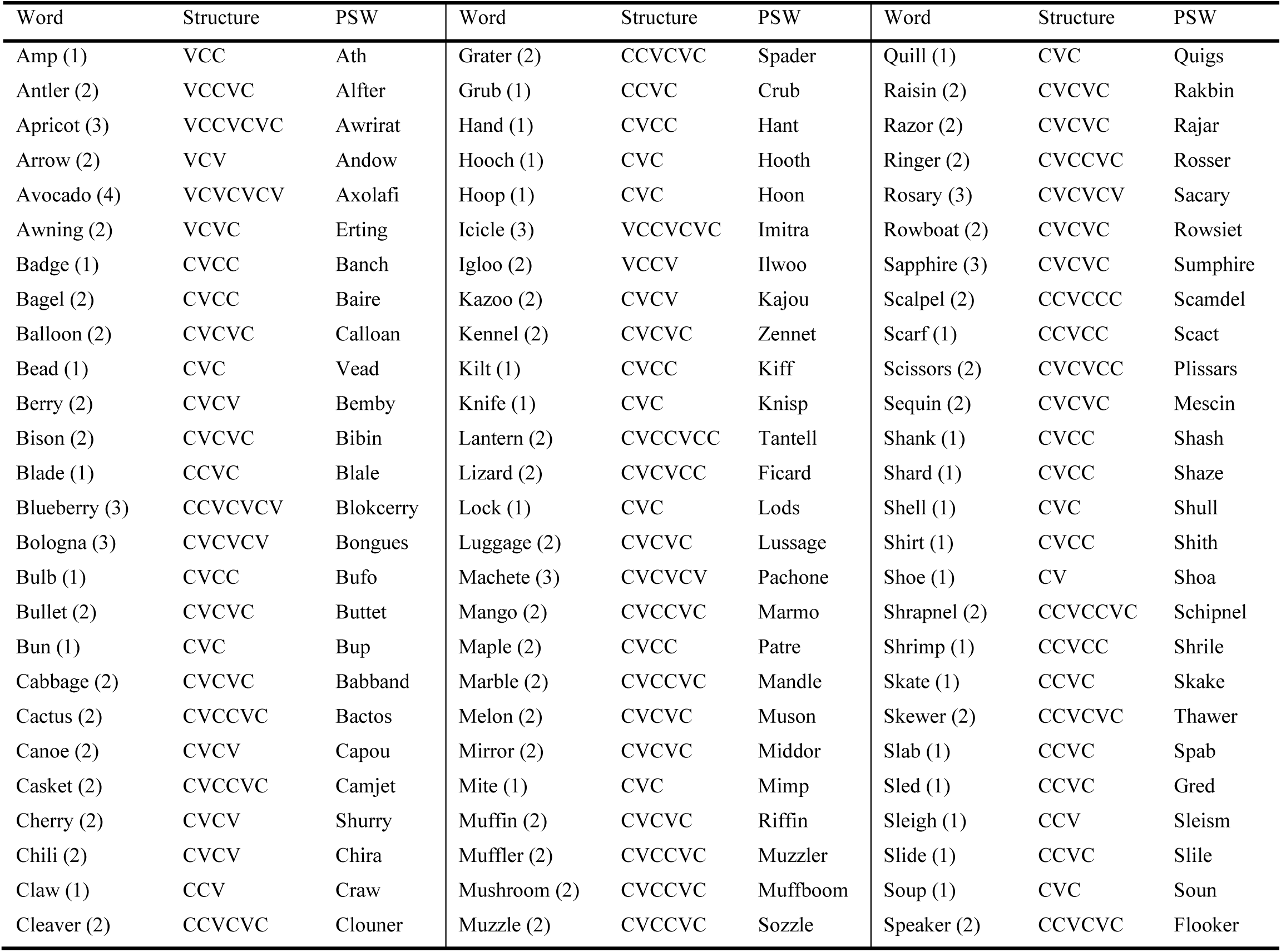
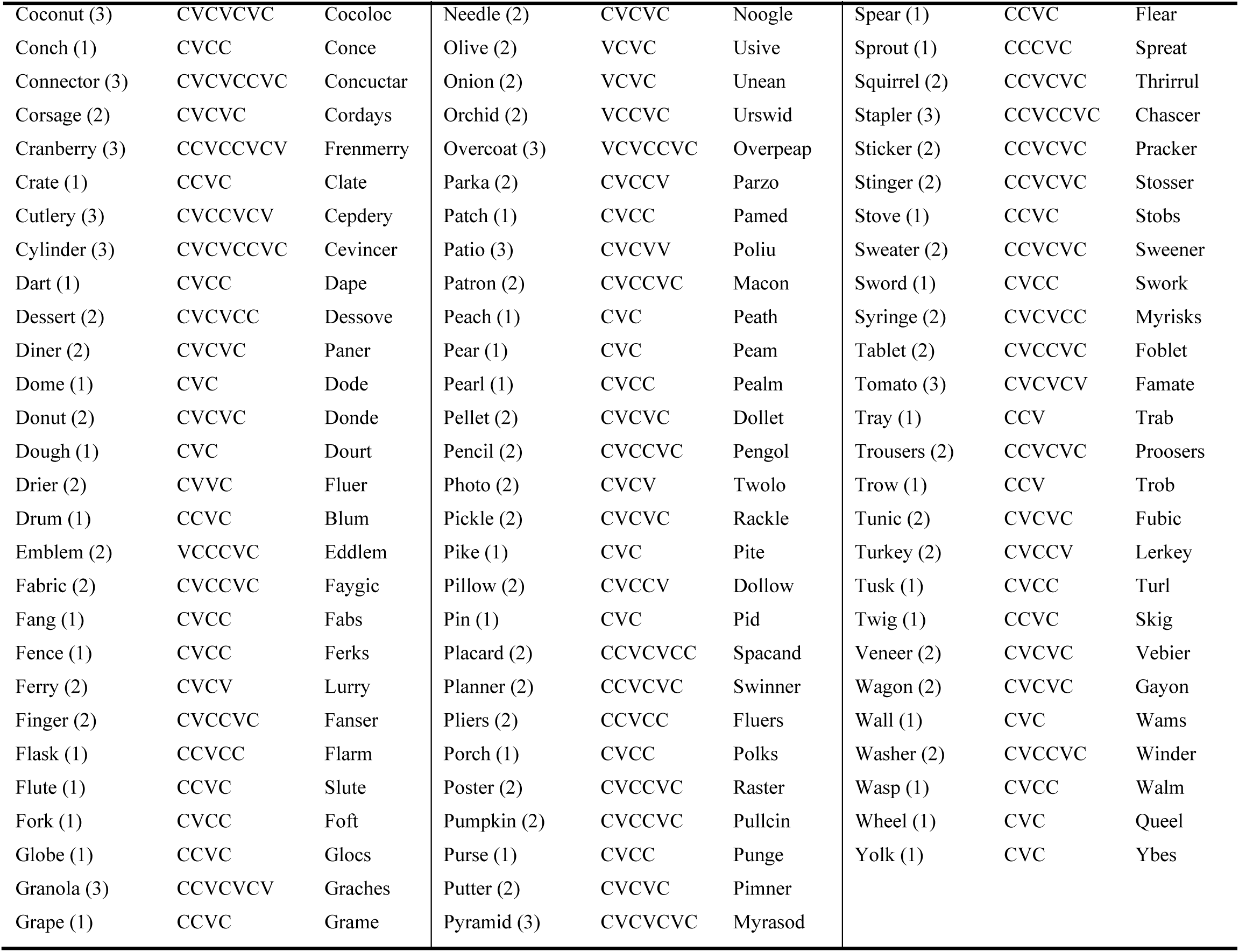
Real words, their syllables, consonant-vowel structure, and corresponding derived pseudoword.

### CVCV pseudowords

We used 320 CVCV pseudowords (duration: *M* = 635 ms, *S* = 104 ms). These were a subset of the 537 pseudowords used by Lacey et al. (2020), for which McCormick et al. (2015) had previously obtained ratings of how rounded/pointed each pseudoword sounded, using seven-point rating scales. The 320 CVCV pseudowords used in the present work were selected based on the prior ratings. These pseudowords were chosen to form a distribution from very rounded (n = 40) to very pointed (n = 40) ratings with the remaining items extending evenly away from either side of the median forming slightly rounded (n = 120) and slightly pointed (n = 120) groups. As elaborated on in the Supplemental Methods, these pseudowords were selected through an iterative process; beginning with items closest to the endpoints/median, items were replaced to ensure the final set had a relatively uniform biphone frequency.

### Derived pseudowords

160 pseudowords (duration: *M* = 626 ms, *S* = 121 ms) were generated by replacing parts of the real words, making these pseudowords more phonologically similar to the real words than the CVCV pseudowords. In contrast to the CVCV pseudowords, these were neither strictly CVCV in structure nor strictly disyllabic. They were generated using the software "Wuggy" (Keuleers & Brysbaert, 2010), which alters the spelling of real words to make phonotactically similar pseudowords. One such pseudoword was selected for each real word. As Wuggy generates orthographic pseudowords, only pronounceable pseudowords were selected. A complete list of the derived pseudowords and the corresponding real word from which each was derived is provided in Table 1.

### Acoustic parameters

Lacey et al. (2020) found correlations between shape ratings and six acoustic measures of voice quality and regularity (vocal parameters): the fraction of unvoiced frames (FUF), mean autocorrelation, pulse number, jitter, shimmer, and harmonics-to-noise ratio (HNR). Among these parameters FUF, mean autocorrelation, pulse number, and HNR all measure the relative amount of voiced and unvoiced information in an utterance; while shimmer and jitter provide information about the amount of acoustic variation in an utterance. In more detail, the FUF is the percentage of time windows with voicing present across the duration of the speech item (Boersma & Weenink, 2012). Mean autocorrelation is the correlation of the acoustic signal between time-shifted versions of itself (Boersma & Weenink, 2012). Pulse number is the number of glottal openings and closings over the duration of a speech item (Boersma & Weenink, 2012). Jitter and shimmer are measures of glottal source variability, captured by the variability in the frequency and amplitude of the speech acoustics, respectively (Teixeira & Fernandes, 2014). The HNR is the ratio of periodic to aperiodic proportions of a speech item’s acoustic signal (Teixeira & Fernandes, 2014). Kumar et al. (2025) found that shape ratings also significantly correlated with the standard deviation of the fundamental frequency (F0 SD) and the duration of the pseudoword, but not the mean fundamental frequency (mean F0). Here, we extracted all nine of these vocal parameters (a single value for each per item) with custom Praat code (Boersma & Weenink, 2011).

Lacey et al. (2020) also used representational similarity analyses to show that perceptual shape ratings of pseudowords were associated with three spectro-temporal parameters: spectral tilt (the slope of the power spectral density; Aiken & Picton, 2008; Oppenheim, 1970), the speech envelope (amplitude profile), and the FFT (the power at different frequencies; see also Kumar et al., 2025, for a replication). As in Lacey et al. (2020), for this set of stimuli, each spectro-temporal parameter was measured with multiple values per item, extracted using custom MATLAB code (Mathworks Inc. version 2020a; The MathWorks, Natick, MA). We first normalized the duration of the stimuli to the mean duration for their respective set by removing or interpolating data points using the resampling function in MATLAB (Lacey et al., 2020). This ensured that the calculations done on these parameters were based on the same number of values for each item in each set (although, due to the different calculations involved for each spectro-temporal parameter, this number differed across parameters; see Table S1). From these measures, we calculated the coefficient of variation (standard deviation/mean) for each spectro-temporal parameter in each stimulus set. The coefficient of variation is a metric of variability and thus could capture relationships between variation in the values of the acoustic parameters and shape ratings.

### Perceptual rating task

The experiment was coded using PsychoPy (Peirce, 2007) and run online at https://pavlovia.org/. All participants were asked to rate how rounded or pointed the real words, derived pseudowords, and CVCV pseudowords sounded. The three stimulus types were presented to each participant in separate blocks. The order of blocks was counterbalanced across participants. Each block presented every stimulus item once, and the order of items was randomized for each participant. On-screen instructions at the start of the experiment explained the rating task to each participant. Participants were randomly assigned to rate the sound of each item using either a rounded-to-pointed (1 = "very rounded"; 7 = "very pointed") or pointed-to-rounded (1 = "very pointed"; 7 = "very rounded") numeric scale. Rating responses were entered using the number keys at the top of the keyboard.

## RESULTS

### Iconicity associations of current stimuli

A necessary condition for assessing iconicity is that item ratings are systematic across participants. We began by evaluating the reliability of item ratings using a series of Cronbach’s α coefficients, computed separately for each stimulus set and rating scale (rounded-to-pointed vs. pointed-to-rounded).

The real words showed robust reliability, with α = 0.83 and α = 0.77 for the rounded-to-pointed and pointed-to-rounded scales, respectively. This indicates a high level of consistency as would be expected for iconic stimuli. The CVCV pseudowords were found to have strong reliability (α = 0.80) for the rounded-to-pointed scale and weaker but still acceptable reliability (α = 0.51) for the pointed-to-rounded scale, indicating that the ratings of these pseudowords are also systematic. Finally, we consider the derived pseudowords, which unlike the CVCV pseudowords were not designed to systematically explore iconic sound to meaning correspondences, but were instead designed to be structurally word-like and matched to the real words. These ratings were also very reliable, with α = 0.83 and α = 0.70 for the rounded-to-pointed and pointed-to-rounded scales, respectively.

### Comparing the acoustic-to-rating relationship across real words and pseudowords

Our primary analyses tested whether pseudowords and real words have similar acoustic-to-rating relationships. This comparison tests whether lexical status (real word vs. pseudoword) influences iconicity. To simplify these analyses, we combined ratings from the rounded-to-pointed and pointed-to-rounded scales (see Lacey et al., 2020). Before rescaling, we tested the correlations between opposing scales (see Lacey et al., 2020). The negative correlations between scales were significant for the real words (*r_158_* = -0.82, *p* < .001), CVCV pseudowords (*r_318_* = - 0.67, *p* < .001), and derived pseudowords (*r_158_* = -0.73, *p* < .001). These results indicate that, for all stimulus sets, the two scales can be considered to measure the same construct. Moreover, these results support our decision to re-code the ratings to a single scale for subsequent testing.

It is possible that the ratings of real words could be biased by participants’ knowledge of their meanings. To evaluate whether real word ratings were driven by sound rather than (or in addition to) semantics, we correlated the mean ratings for real words with those for their corresponding derived pseudowords. This correlation was significant (*r*_158_ = .35, *p* < .001), indicating that the real words and their derived pseudowords tended to have similar iconic associations and that the real word ratings were at least partly based on their sounds. To better understand the relationship between the ratings for the real words, their sound, and their meaning, we next conducted an exploratory analysis that compared our data to the sound scores and meaning scores reported by Sidhu et al., (2021). To this end, we re-scaled our ratings to correspond to the pointed to rounded scale used by Sidhu et al., (2021). The real word ratings were strongly correlated with the sound score, *r_158_* = .63, *p <* .001, suggesting that the real word ratings reflected their sound. There was also a strong correlation between ratings and the meaning score, *r_158_* = .66, *p <* .001, which is expected given the strong phoneme-to-meaning relationship reported by Sidhu et al., (2021). Importantly, a partial correlation between real word ratings and the ratings for their corresponding derived pseudowords while controlling for meaning was significant (*r_158_* = .25, *p <* .001), further supporting the contention that ratings of the real words were, at least partially, driven by their sounds rather than their semantics.

The mean rounded-to-pointed ratings for each stimulus were next entered into separate Pearson correlations for each acoustic parameter and stimulus set. The alpha level for these correlations was set by Bonferroni correction for 12 comparisons (α = .0042). Figures 1–3 show the correlations between mean ratings (on the rounded-to-pointed scale) and each acoustic parameter for the real words, derived pseudowords, and CVCV pseudowords. Although the exact values differ across sets, the overall patterns and rankings of correlations are notably similar, a pattern that is illustrated quite clearly in Figure 4. This suggests that the acoustic-to-rating relationships observed for pseudowords are also present for real words.

**Figure 1.**
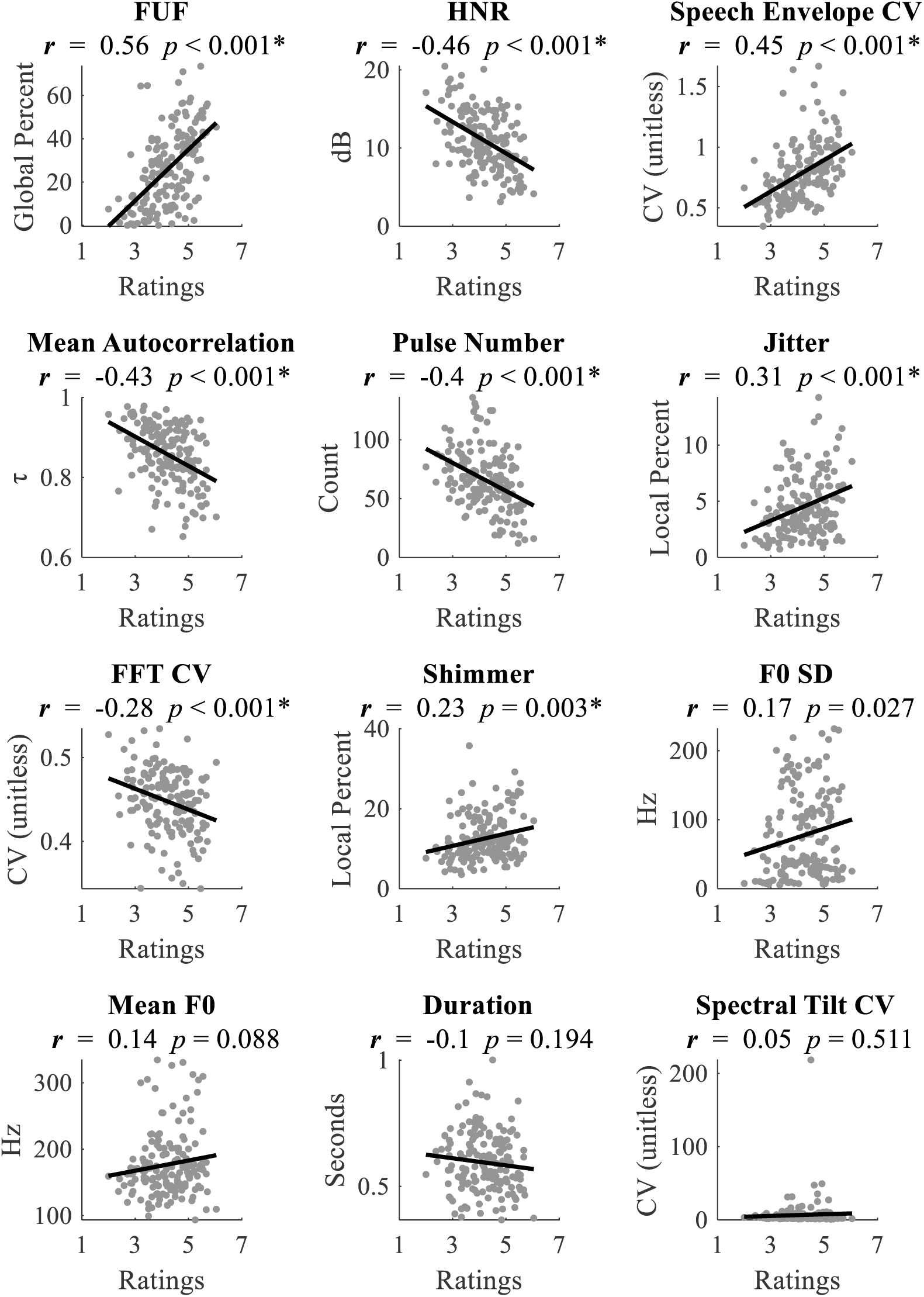
Pearson correlations between the mean ratings (horizontal axes; 1-7 based on the limits of the “very rounded” to “very pointed” scale) and each acoustic parameter (vertical axes) for the 160 real words. Ordered by correlation size. Bonferroni corrected α for twelve correlations is *p* < .0042; * indicates significant after correction. Abbreviations: CV: coefficient of variation; FFT: fast Fourier transform; FUF: fraction of unvoiced frames; F0: fundamental frequency; HNR: harmonics to noise ratio.

**Figure 2.**
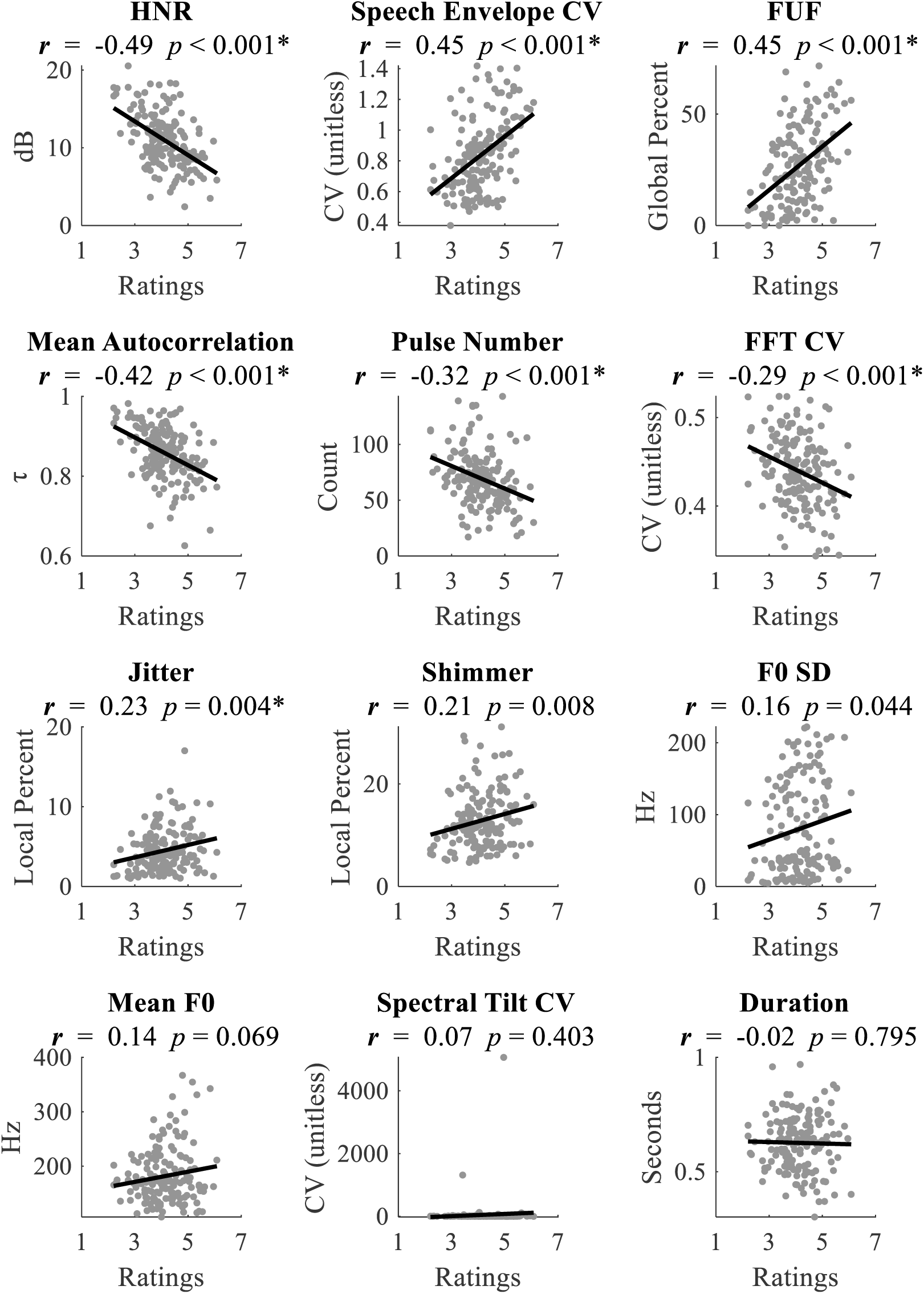
Pearson correlations between the mean ratings (horizontal axes; 1-7 based on the limits of the “very rounded” to “very pointed” scale) and each acoustic parameter (vertical axes) for the 160 derived pseudowords. Ordered by correlation size. Bonferroni corrected α for twelve correlations is *p* < .0042; * indicates significant after correction. Abbreviations as in Figure 1.

**Figure 3.**
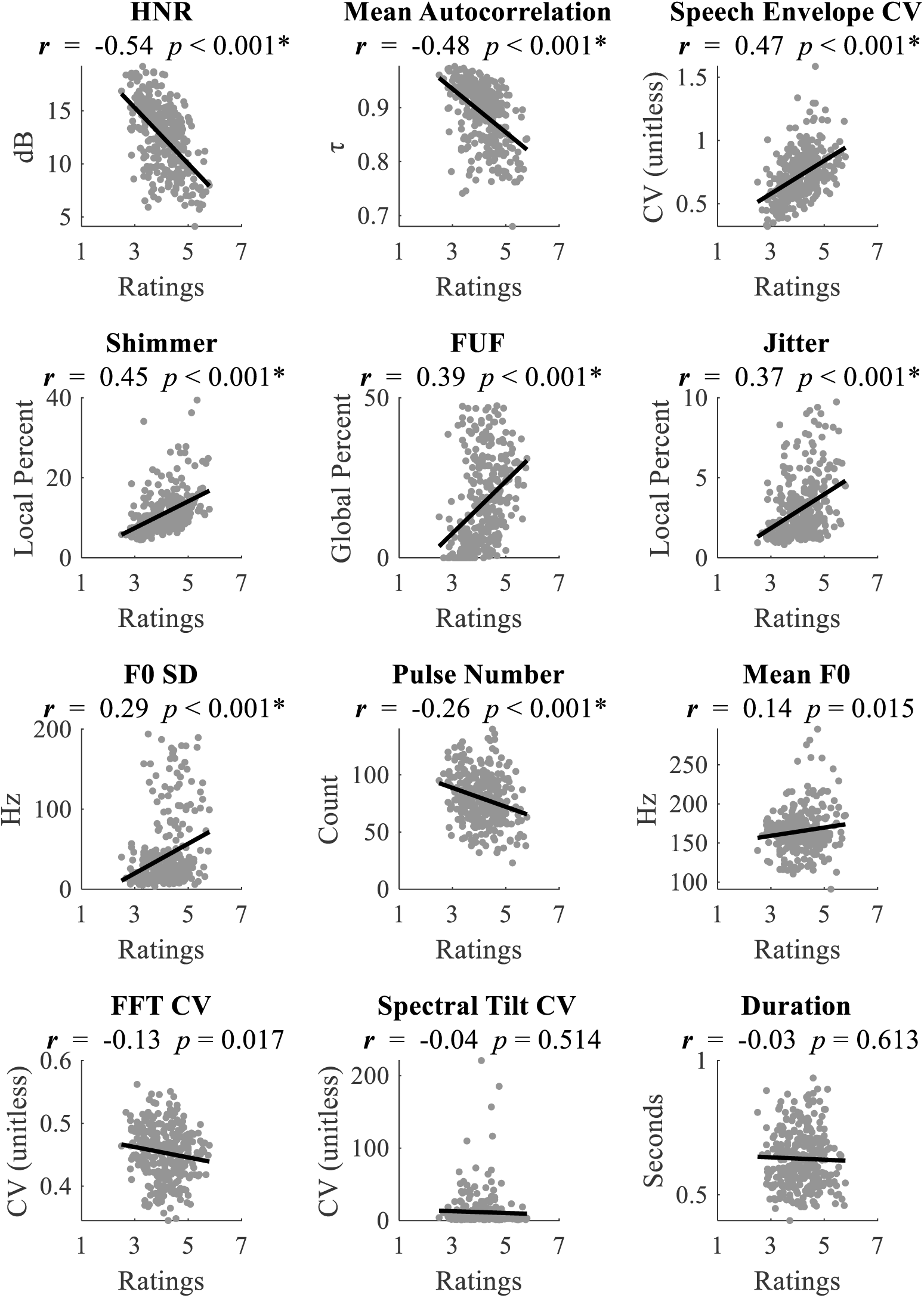
Pearson correlations between the mean ratings (horizontal axes; 1-7 based on the limits of the “very rounded” to “very pointed” scale) and each acoustic parameter (vertical axes) for the 320 CVCV pseudowords. Ordered by absolute value of correlation. Bonferroni corrected α for twelve correlations is *p* < .0042; * indicates significant after correction. Abbreviations as in Figure 1.

**Figure 4.**
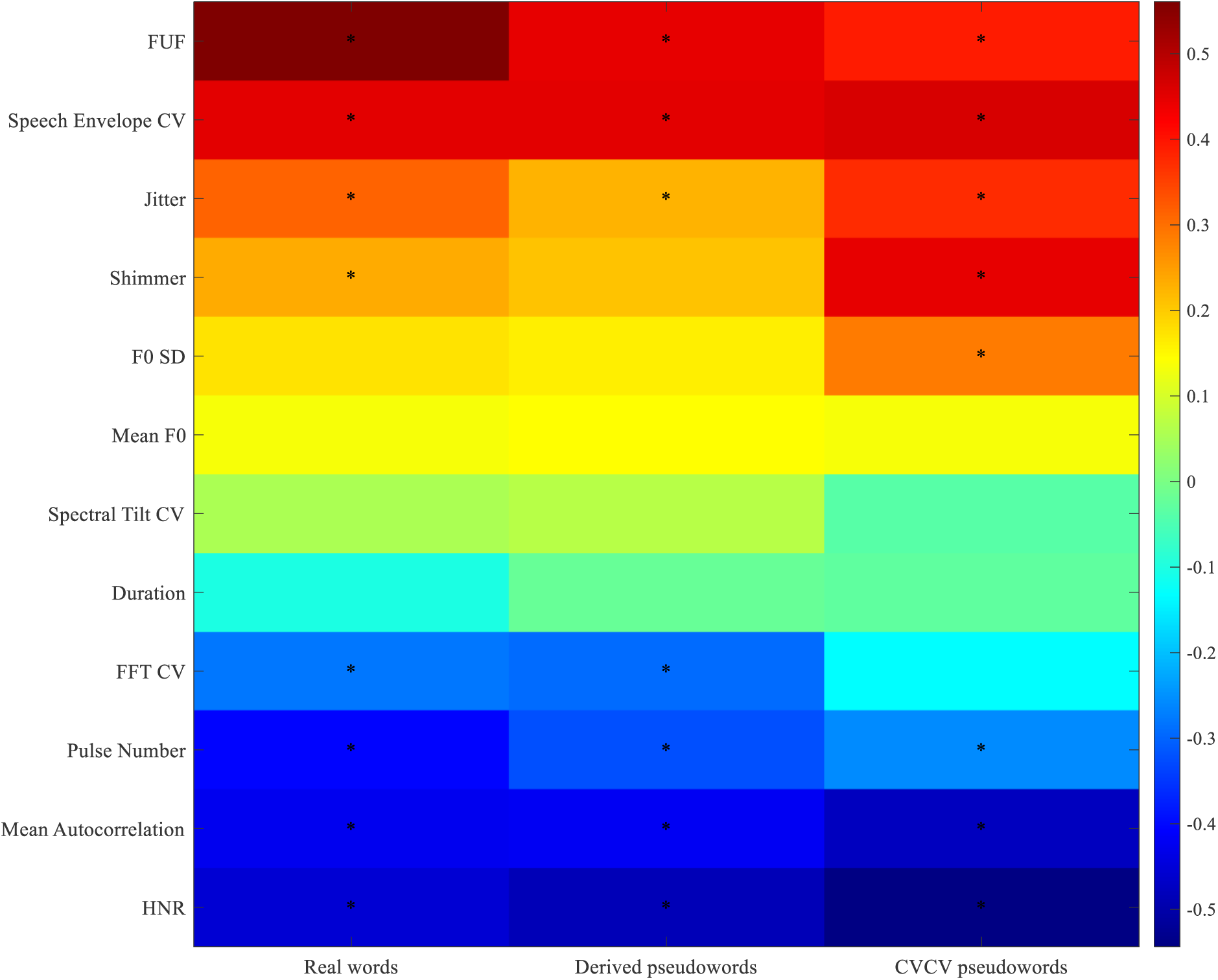
The ratings-to-acoustic parameter correlations for real words, derived pseudowords, and CVCV pseudowords. Plotted values are the Pearson correlation coefficients reported in figures 1-3; * mark significant correlations as noted in those figures.

## DISCUSSION

This study tested whether the relationship between the acoustic properties of spoken pseudowords and judgments of shape iconicity (Lacey et al., 2020; Kumar et al., 2025; Nayak et al., 2024) generalizes to spoken real words. We collected ratings for 160 real words and 160 word-like pseudowords derived from those real words. We also collected shape ratings for a 320-item subset of the CVCV pseudowords used in earlier work (Lacey et al., 2020; Kumar et al., 2025). We measured the acoustic-to-rating relationship through correlations between ratings and 12 acoustic features. Finally, to test if shape iconicity is comparable across real words and pseudowords, we compared these acoustic-to-rating correlations between real words and pseudowords. Critically, we found that pseudowords and real words showed similar iconic acoustic-to-rating relationships.

### Speech acoustics in shape iconicity

The results indicate that acoustic-to-rating mappings in pseudowords extend to real words. Briefly, for all three stimulus types, as item ratings changed from rounded to pointed, values of the FUF, jitter, and the coefficient of variation for the speech envelope all increased, and the values of the HNR, mean autocorrelation, and pulse number decreased. The HNR, mean autocorrelation, pulse number, and the FUF are measures of periodicity, and the correlations of these parameters with ratings indicate that less overall periodicity was associated with more pointed judgments. The correlations of ratings with the mean autocorrelation, jitter, and the coefficient of variation for the speech envelope could indicate that stable (or low variance) speech acoustics are associated more with roundedness than pointedness. While only significant for real words and CVCV pseudowords (but marginally significant for derived pseudowords), the correlations of ratings with shimmer also fit with this interpretation. It is also worth noting that the correlations with the FUF and pulse number could indicate a role of voicing in shape iconicity judgments, consistent with prior findings (see Cuskley et al., 2017; D’Onofrio, 2014; Lacey et al., 2026; McCormick et al., 2015).

Correlations with other parameters were less consistent and thus are interpreted more cautiously. Duration, mean F0, and the coefficient of variation for the spectral tilt were not significantly correlated with shape ratings for any stimulus set. The coefficient of variation for the FFT did not significantly correlate with ratings of CVCV pseudowords, but significant correlations for real word and derived pseudoword ratings indicate that higher values of the coefficient of variation for the FFT (i.e., more variable spectral power) corresponded to more rounded ratings. This contrasts with our results for mean autocorrelation, jitter, and coefficient of variation for the speech envelope which indicate that acoustic variability corresponds to more pointed ratings. The explanation for this discrepancy is not clear. The FFT is a high dimensional parameter and it is possible that compressing it into the coefficient of variation, while an essential data reduction for the current work, obscures the true relationship between the FFT and shape ratings.

F0 SD did not significantly correlate with ratings for real words or derived pseudowords, but a significant correlation for CVCV pseudowords indicates that items with higher F0 SD were rated as more pointed. Finally, while shimmer did not significantly correlate with ratings for derived pseudowords, significant correlations for real words and CVCV pseudowords indicate that items with higher shimmer values were rated as more pointed. The F0 SD and shimmer correlations fit the general relationship of more pointed ratings being associated with speech items that exhibit more acoustic variability as noted in the previous paragraph.

### Linking iconicity of pseudowords and real words

A key complication to understanding the relationship between word sounds and meaning is that knowledge of a word’s meaning can bias judgments of how it sounds. To avoid this confound, many investigations rely on pseudowords, implicitly assuming that iconic relationships observed in pseudowords will generalize to real words. Sidhu et al. (2021) demonstrated a phoneme-to-meaning relationship, but because their judgments were made on written forms, they could not isolate the contribution of speech acoustics. By comparing the iconicity ratings to acoustic relationships of auditory pseudowords and real words, the present work provides a direct comparison of the relationship between speech sounds and meaning across lexical status. We report that real words and pseudowords show similar acoustic-to-rating relationships, offering clear evidence that iconic acoustic patterns extend to real spoken words.

### Generalizability

By focusing on speech acoustics, the present results are more directly applicable to spoken language processing than are studies relying on orthographic stimuli. As noted above, rounded/pointed judgments of speech correspond to acoustic factors such as the F2 and F3 frequency (Knoeferle et al., 2017). Recent work examining the basis of iconicity for auditory pseudowords has identified acoustic features associated with roundedness and pointedness judgments (e.g., Lacey et al., 2020; Kumar et al., 2025). Our CVCV pseudowords were a subset of the stimuli reported on by Lacey et al. (2020; Kumar et al., 2025), and produce markedly similar results as those reported previously (see Supplemental Analyses). Though not central to the aims of this investigation the similarity of results for these two stimulus sets demonstrates a cross-talker generalization of shape ratings for phonetically matched pseudowords. Future work should expand on this.

### Conclusion

This work systematically tests whether the speech acoustics associated with iconic shape ratings of pseudowords generalize to real words. This converges with prior findings concerning speech acoustics and iconicity (e.g., Lacey et al., 2020). Across multiple tests, it is apparent that the acoustic-to-rating relationship is substantially similar for real words and pseudowords, offering evidence of iconicity in spoken language. This work addresses a key question concerning our understanding of iconicity in natural language.

## Supporting information

Supplemental Methods, Results, & Tables

## ACKNOWLEDGMENTS

An AI-based language tool assisted in this work. Portions of the text were edited for clarity and style by Microsoft Copilot. The authors take full responsibility for the content. An AI-based language tool (Microsoft Copilot) produced portions of the code and command syntax for generating figures and handling data. All code was reviewed, validated, and executed by the author, who confirmed the accuracy of each step and the integrity of the resulting data and figures.

## Declarations

### Ethics approval

The institutional review board of Penn State College of Medicine approved all procedures for the reported research.

### Consent to participate

Consent was obtained from all participants prior to data collection.

### Consent for publication

All authors approve submissions of this work to this journal.

### Availability of data and materials

Data and relevant materials are available upon request.

### Code availability

Relevant code will be made available upon request.

### Authors contributions

JD, SL, LCN, KS designed the research. JD collected the data. JD analyzed the data. JD, SL, LCN, and KS wrote the paper. All authors contributed to study design, interpretation of results, and manuscript revision, and approved the final version.

## Supplemental Materials

### Method

#### Selecting real words

Using data reported by Sidhu et al. (2021), we ensured that our selected words did not differ in concreteness (*F_3,156_* = 2.31, *p* = .078) or lexical frequency (*F_3,156_* = 0.66, *p* = 0.58). Sidhu et al. (2021) provide estimates of word concreteness and lexical frequency, which we used to evaluate our selected words. Concreteness refers to how abstract a word’s meaning is and is often derived from participant ratings on abstract-to-concrete rating scales. Lexical frequency is a measure of how often a word occurs in a language (Brysbaert & New, 2009). Next, using data from the Irvine Phonotactic Dictionary (IPhoD; Vaden et al., 2009), we ensured that our selected items did not differ in similar number of syllables (*F_3,156_* = 0.63, *p* = 0.60), stressed phonological neighborhood density (*F_3,156_* = 0.36, *p* = 0.78), and biphone probability (*F_3,156_* = 0.39, *p* = 0.76). Phonological neighborhood density is the number of words that could be made by changing a single phoneme in a word, and biphone probability is the probability of two phonemes being adjacent in English. These word characteristics are further summarized in Table S2, along with Sidhu et al. (2021) measures of word rounded/pointed meaning, rounded/pointed sound, and “Derived Iconicity” (the agreement between a word’s meaning and sound).

#### Selecting CVCV pseudowords

We used the IPhoD calculator (Vaden et al., 2009) to calculate the biphone probability of each pseudoword. We ensured that the selected 40 extreme rounded and 40 extreme pointed pseudowords (based on the McCormick et al. 2015 ratings) and the 240 near median pseudowords (divided into 40 item batches) did not differ in biphone frequency (*F_7_,*_312_ = .84, *p* = 0.56).

### Analyses

#### Comparison to prior findings

As noted in the introduction section, recent work (Lacey et al., 2020; Kumar et al., 2025; Nayak et al., 2024) reports reliable correlations between shape ratings and acoustic parameters in pseudowords. While using many of the same pseudowords, the stimuli from this study were produced by a different talker than the Lacey et al. (and Kumar et al., 2025) items. Moreover, the current experiment collected shape ratings using rounded-to-pointed/pointed-to-rounded scales, while the prior work used "not rounded" to "very rounded"/"not pointed" to "very pointed" scales. These methodological differences should not influence the relationship between shape rating and acoustic parameters reported previously (Lacey et al., 2020; Kumar et al., 2025). Nonetheless, the 320 bi-syllable pseudowords included in this experiment allow us to test this assumption. If the ratings with these pseudowords are consistent with the results reported by Lacey et al. (2020; Kumar et al., 2025), then the methodological difference between these studies is unlikely to be a factor in our analyses using the real word and derived pseudoword stimuli.

##### Item-wise ratings

As a first step, we tested how similar the ratings of the current CVCV pseudowords were to their ratings obtained when they were part of the more extensive set used in McCormick et al. (2015). The mean rounded-to-pointed rating for our stimuli was strongly correlated with their mean rating from both McCormick et al. (2015), *r_318_* = 0.61, *p* < .001, and Kumar et al. (2025), *r_318_* = 0.65, *p* < .001. Thus, our stimuli and ratings procedure produce ratings consistent with previously reported item-wise ratings.

##### Vocal parameter correlations

Next, we tested the correlations between the mean rounded-to-pointed ratings for the CVCV pseudowords (see main text) and the nine vocal parameters noted in prior work (e.g., Lacey et al., 2020; Kumar et al., 2025). We include the correlations reported in that prior work in Table S3. Briefly, these correlations show that pseudowords with a lower FUF, lower jitter, and lower shimmer were associated with more rounded ratings, while pseudowords with a higher HNR, a higher number of pulses, and higher mean autocorrelation tended to be rated as sounding more rounded.

All vocal parameters, except duration and mean F0, significantly correlated with the mean ratings. The pattern of significant effects in the current data broadly overlaps with those reported in prior work, with most parameters that reached significance here also reported as significant in at least one previous study (Table S3). Like the current results, the correlation with mean F0 was consistently non-significant across prior work. In addition to the similar pattern of statistical significance, these correlations were all in the same direction and were mainly of the same magnitude as those reported by Lacey et al. (2020; Kumar et al., 2025). In addition to Lacey et al. (2020; Kumar et al., 2025), another recent study (Nayak, 2024) also tested the correlation between these vocal parameters and the rounded-to-pointed ratings, albeit with a different set of CVCV pseudowords (Table S2). Despite these differences, the present set of correlations is notably similar to what is reported by Nayak (2024). Thus, the current findings are consistent with the correlations between vocal parameters and mean ratings reported by three earlier studies using two different CVCV pseudoword sets.

## Tables

**Table S1.**
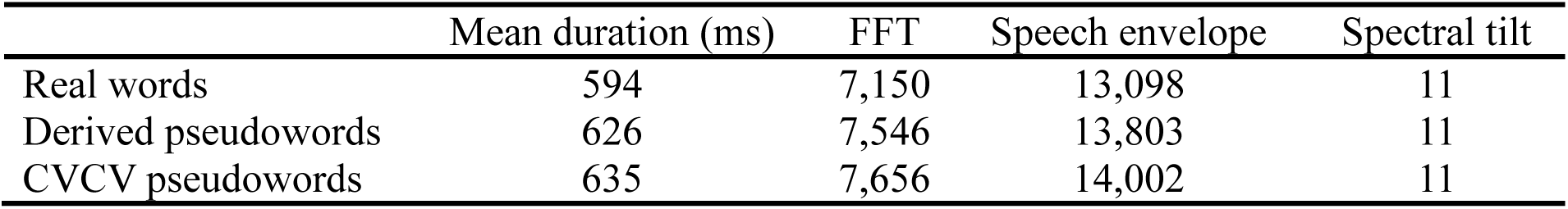
Item duration and normalized spectral-temporal vector lengths.

**Table S2.**
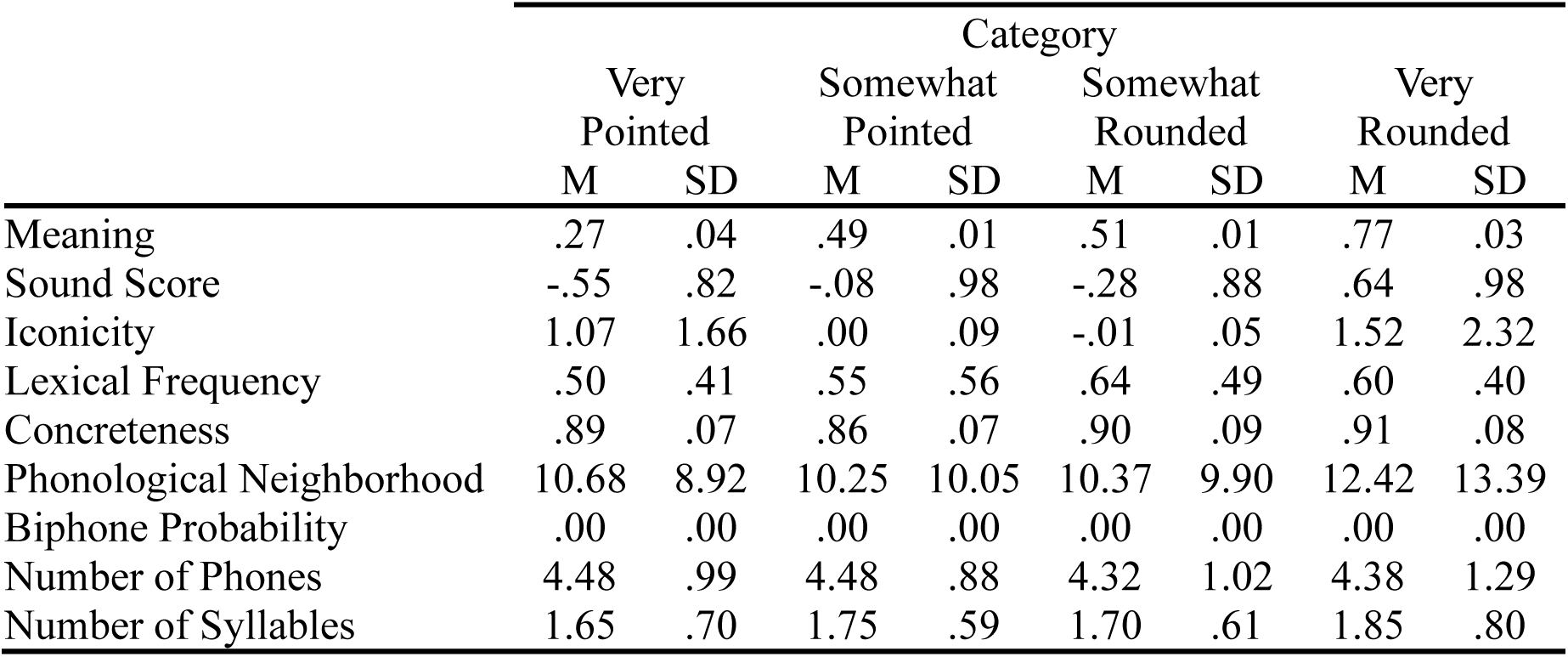
Characteristics of selected words.

**Table S3.**
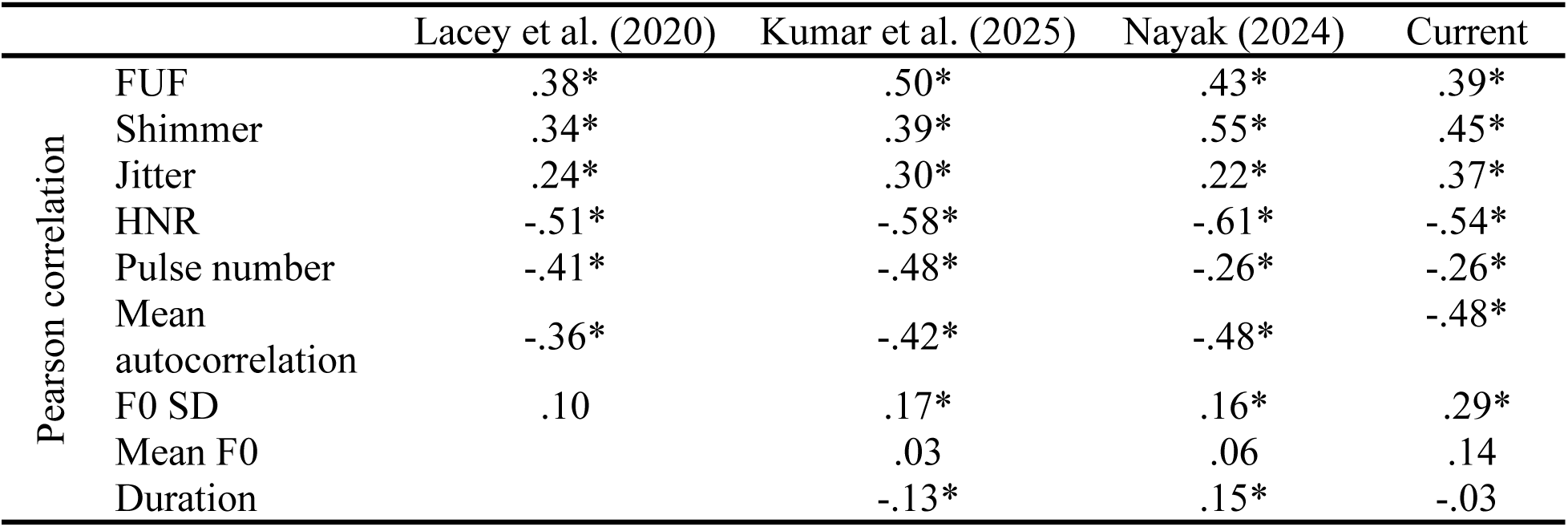
Acoustic-to-ratings correlations for CVCV pseudowords in prior and current work. *statistically significant.

## Footnotes

1 Vowel formants refer to the resonance frequencies of the human vocal tract. The first three formants, F1-F3, are primarily informative about vowel identity while the other formants relate more to speaker identity (Knoeferle et al., 2017).

## REFERENCES

Ademollo, F. (2011). The Cratylus of Plato: A Commentary. Cambridge University Press: 3 Cambridge, UK.

Ahlner, F., & Zlatev, J. (2010). Cross-modal iconicity: A cognitive semiotic approach to sound symbolism. Sign Systems Studies, 38(1/4), 298–348. 10.12697/sss.2010.38.1-4.11

Aiken, S. J., & Picton, T. W. (2008). Envelope and spectral frequency-following responses to vowel sounds. Hearing Research, 245(1–2), 35–47. 10.1016/j.heares.2008.08.004

Akita, K., & Tsujimura, N. (2016). Handbook of Japanese Lexicon and Word Formation. In T. Kageyama & H. Kishimoto (Eds.), Handbook of Japanese Lexicon and Word Formation (Eds.). Walter de Gruyter GmbH & Co KG. 10.1515/9781614512097

Blasi, D. E., Wichmann, S., Hammarström, H., Stadler, P. F., & Christiansen, M. H. (2016). Sound–meaning association biases evidenced across thousands of languages. Proceedings of the National Academy of Sciences, 113, 10818–10823.

Boersma, P., & Weenink, D. (2011). Praat: Doing Phonetics by Computer. Ear & Hearing, 32(2), 266. 10.1097/aud.0b013e31821473f7 https://doi.org/10.3758/BRM.41.4.977

Catricalà, M., & Guidi, A. (2015). Onomatopoeias: a new perspective around space, image schemas and phoneme clusters. Cognitive Processing, 16, 175–178. 10.1007/s10339-015-0693-x

Cuskley, C., Simner, J., & Kirby, S. (2017). Phonological and orthographic influences in the bouba–kiki effect. Psychological Research, 81(1), 119–130. 10.1007/s00426-015-0709-2

D’Onofrio, A. (2014). Phonetic Detail and Dimensionality in Sound-shape Correspondences: Refining the Bouba-Kiki Paradigm. Language and Speech, 57(3), 367–393. 10.1177/0023830913507694

Greenberg, J. H., & Jenkins, J. J. (1966). Studies in the Psychological Correlates of the Sound System of American English. WORD, 22(1–3), 207–242. 10.1080/00437956.1966.11435451

Hockett, C. F. (1960). The Origin of Speech. Scientific American, 203(3), 88–97.

IBM Corp. (2022). IBM SPSS Statistics for Windows (Version 29.0) [Computer software]. IBM Corp

Keuleers, E., & Brysbaert, M. (2010). Wuggy: A multilingual pseudoword generator. Behavior Research Methods, 42(3), 627–633. 10.3758/BRM.42.3.627

Knoeferle, K., Li, J., Maggioni, E., & Spence, C. (2017). What drives sound symbolismDifferent acoustic cues underlie sound-size and sound-shape mappings. *Scientific Reports*, December 2016, 1–11. 10.1038/s41598-017-05965-y

Köhler, W. (1929). Gestalt Psychology. New York: Liveright Publishing Corporation.

Köhler, W. (1947). Gestalt Psychology: An Introduction to New Concepts in Modern Psychology. Liveright: New York, NY

Kumar, G. V., Lacey, S., Dorsi, J., Nygaard, L. C., & Sathian, K. (2025). Acoustic parameter combinations underlying mapping of pseudoword sounds to multiple domains of meaning: Representational similarity analyses and machine-learning models. The Journal of the Acoustical Society of America, 158(6), 4243–4267

Lacey, S., Jamal, Y., List, S. M., McCormick, K., Sathian, K., & Nygaard, L. C. (2020). Stimulus parameters underlying sound-symbolic mapping of auditory pseudowords to visual shapes. Cognitive Science, 44(e12883). 10.1111/cogs.12883

Lacey, S., Matthews, K., Shrivastava, A., Sathian, K. & Nygaard, L.C. (2026). Domain-specificity in phonological and acoustic characteristics of pseudoword iconicity across multiple meaning domains. (Preprint, bioRxiv, doi: 10.1101/2024.09.03.610973).

Lockwood, G., & Dingemanse, M. (2015). Iconicity in the lab: a review of behavioral, developmental, and neuroimaging research into sound-symbolism. Frontiers in Psychology, 6(August), 1–14. 10.3389/fpsyg.2015.01246

Lockwood, G., Hagoort, P., & Dingemanse, M. (2016). How Iconicity Helps People Learn New Words : Neural Correlates and Individual Differences in Sound-Symbolic Bootstrapping. 2(1), 1–15.

Maurer, D., Pathman, T., & Mondloch, C. J. (2006). The shape of boubas: Sound-shape correspondences in toddlers and adults. Developmental Science, 9(3), 316–322. 10.1111/j.1467-7687.2006.00495.x

Mazzoni, D. (2006). Audacity. http://audacity.sourceforge.net/

McCormick, K., Kim, J. Y., List, S., & Nygaard, L. C. (2015). Sound to meaning mappings in the bouba-kiki effect. In D. C. Noelle, R. Dale, A. S. Warlaumont, J. Yoshimi, T. Matlock, C. D. Jennings, & P. P. Maglio (Eds.), Proceedings 37th Annual Meeting Cognitive Science Society (pp. 1565–1570). Austin TX, USA: Cognitive Science Society.

Miller, G. A., Heise, G. A., & Lighten, W. (1951). The intelligibility of speech as a function of the context of the test materials. The Journal of Experimental Psychology, 41(5), 329–335

Monaghan, P., Shillcock, R. C., Christiansen, M. H., & Kirby, S. (2014). How arbitrary is language? Philosophical Transactions of the Royal Society B: Biological Sciences, 369(1651). 10.1098/rstb.2013.0299

Nayak, S. (2024). Phonetic and acoustic analyses of sound-symbolic associations of a large pseudoword set across multiple domains of meaning. Unpublished master’s thesis, Pennsylvania State University, College of Medicine, Hershey, PA, USA.

Newman, S. S. (1933). Further experiments in phonetic symbolism. The American Journal of Psychology, 45(1), 53. 10.2307/1414186

Nygaard, L. C., Cook, A. E., & Namy, L. L. (2009). Sound to meaning correspondences facilitate word learning. Cognition, 112(1), 181–186. 10.1016/j.cognition.2009.04.001

Oppenheim, A. V. (1970). Speech spectrograms using the fast Fourier transform. IEEE Spectrum, 7(8), 57–62. 10.1109/mspec.1970.5213512

Parise, C. V., & Pavani, F. (2011). Evidence of sound symbolism in simple vocalizations. Experimental Brain Research, 214(3), 373–380. 10.1007/s00221-011-2836-3

Peirce, J. W. (2007). PsychoPy-Psychophysics software in Python. Journal of Neuroscience Methods, 162(1–2), 8–13. 10.1016/j.jneumeth.2006.11.017

Ramachandran, V. S., & Hubbard, E. M. (2001). Synaesthesia - A window into perception, thought and language. Journal of Consciousness Studies, 8(12), 3–34.

Sapir, E. (1929). A study in phonetic symbolism. Journal of Experimental Psychology, 12(3), 225–239. 10.1037/h0070931

Rubin, P., Turvey, M. T., & Van Gelder, P. (1976). Initial phonemes are detected faster in spoken words than in spoken nonwords. Perception & Psychophysics, 19(5), 394–398

Saussure, F. de (1916/2009). Course in General Linguistics. Open Court Classics: Peru, IL, USA.

Sidhu, D. M., Westbury, C., Hollis, G., & Pexman, P. M. (2021). Sound symbolism shapes the English language: The maluma/takete effect in English nouns. Psychonomic Bulletin and Review, 28(4), 1390–1398. 10.3758/s13423-021-01883-3

Sidhu, D. M., Williamson, J., Slavova, V., & Pexman, P. M. (2022). An investigation of iconic language development in four datasets. Journal of Child Language, 49(2), 382–396. 10.1017/S0305000921000040

Sučević, J., Savić, A. M., Popović, M. B., Styles, S. J., & Ković, V. (2015). Balloons and bavoons versus spikes and shikes: ERPs reveal shared neural processes for shape-sound-meaning congruence in words, and shape-sound congruence in pseudowords. Brain and Language, 145–146, 11–22. 10.1016/j.bandl.2015.03.011

Teixeira, J. P., & Fernandes, P. O. (2014). Jitter, Shimmer and HNR Classification within Gender, Tones and Vowels in Healthy Voices. Procedia Technology, 16, 1228–1237. 10.1016/j.protcy.2014.10.138

Tzeng, C. Y., Nygaard, L. C., & Namy, L. L. (2017). Developmental change in children’s sensitivity to sound symbolism. Journal of Experimental Child Psychology, 160, 107–118. 10.1016/j.jecp.2017.03.004

Weingartner, R. H. (1970). Making Sense of the “Cratylus.” Phronesis, 15(1), 5–25.

Westbury, C., Hollis, G., Sidhu, D. M., & Pexman, P. M. (2018). Weighing up the evidence for sound symbolism: Distributional properties predict cue strength. Journal of Memory and Language, 99, 122–150. 10.1016/j.jml.2017.09.006

Winter, B., Lupyan, G., Perry, L. K., Dingemanse, M., & Perlman, M. (2024). Iconicity ratings for 14,000+ English words. Behavior Research Methods, 56(3), 1640–1655. 10.3758/s13428-023-02112-6

## Supplemental References

Brysbaert, M., & New, B. (2009). Moving beyond Kučera and Francis: A critical evaluation of current word frequency norms and the introduction of a new and improved word frequency measure for American English. Behavior Research Methods, 41(4), 977–990.

Vaden, K., Halpin, H., & Hickok, G. (2009). Irvine Phonotactic Online Dictionary (p. available from http://www.iphod.com.).

