## Supplemental Methods, Results, & Tables for "Associations Between Shape Iconicity Ratings and Speech Acoustics: Comparing Real Words and Pseudowords"

Table S1
*Item duration and normalized spectral-temporal vector lengths.*

| | Mean duration (ms) | FFT | Speech envelope | Spectral tilt |
|---|---|---|---|---|
| Real words | 594 | 7,150 | 13,098 | 11 |
| Derived pseudowords | 626 | 7,546 | 13,803 | 11 |
| CVCV pseudowords | 635 | 7,656 | 14,002 | 11 |

Table S2
*Characteristics of selected words*

| | Category | | | | | | | |
|---|---|---|---|---|---|---|---|---|
| | Very Pointed | | Somewhat Pointed | | Somewhat Rounded | | Very Rounded | |
| | M | SD | M | SD | M | SD | M | SD |
| Meaning | .27 | .04 | .49 | .01 | .51 | .01 | .77 | .03 |
| Sound Score | -.55 | .82 | -.08 | .98 | -.28 | .88 | .64 | .98 |
| Iconicity | 1.07 | 1.66 | .00 | .09 | -.01 | .05 | 1.52 | 2.32 |
| Lexical Frequency | .50 | .41 | .55 | .56 | .64 | .49 | .60 | .40 |
| Concreteness | .89 | .07 | .86 | .07 | .90 | .09 | .91 | .08 |
| Phonological Neighborhood | 10.68 | 8.92 | 10.25 | 10.05 | 10.37 | 9.90 | 12.42 | 13.39 |
| Biphone Probability | .00 | .00 | .00 | .00 | .00 | .00 | .00 | .00 |
| Number of Phones | 4.48 | .99 | 4.48 | .88 | 4.32 | 1.02 | 4.38 | 1.29 |
| Number of Syllables | 1.65 | .70 | 1.75 | .59 | 1.70 | .61 | 1.85 | .80 |

Table S3
*Acoustic-to-ratings correlations for CVCV pseudowords in prior and current work.*
**statistically significant.*

| | | Lacey et al. (2020) | Kumar et al. (2025) | Nayak (2024) | Current |
|---|---|---|---|---|---|
| Pearson correlation | FUF | .38* | .50* | .43* | .39* |
| | Shimmer | .34* | .39* | .55* | .45* |
| | Jitter | .24* | .30* | .22* | .37* |
| | HNR | -.51* | -.58* | -.61* | -.54* |
| | Pulse number | -.41* | -.48* | -.26* | -.26* |
| | Mean autocorrelation | -.36* | -.42* | -.48* | -.48* |
| | F0 SD | .10 | .17* | .16* | .29* |
| | Mean F0 | | .03 | .06 | .14 |
| | Duration | | -.13* | .15* | -.03 |

### Supplemental References

Brysbaert, M., & New, B. (2009). Moving beyond Kučera and Francis: A critical evaluation of current word frequency norms and the introduction of a new and improved word frequency measure for American English. *Behavior Research Methods*, *41*(4), 977–990.

Kumar, G. V., Lacey, S., Dorsi, J., Nygaard, L. C., & Sathian, K. (2025). Acoustic parameter combinations underlying mapping of pseudoword sounds to multiple domains of meaning: Representational similarity analyses and machine-learning models. *The Journal of the Acoustical Society of America*, *158*(6), 4243-4267

Lacey, S., Jamal, Y., List, S. M., McCormick, K., Sathian, K., & Nygaard, L. C. (2020). Stimulus parameters underlying sound-symbolic mapping of auditory pseudowords to visual shapes. *Cognitive Science*, *44*(e12883). https://doi.org/10.1111/cogs.12883

McCormick, K., Kim, J. Y., List, S., & Nygaard, L. C. (2015). Sound to meaning mappings in the bouba-kiki effect. In D. C. Noelle, R. Dale, A. S. Warlaumont, J. Yoshimi, T. Matlock, C. D. Jennings, & P. P. Maglio (Eds.), *Proceedings 37th Annual Meeting Cognitive Science Society* (pp. 1565–1570). Austin TX, USA: Cognitive Science Society.

Nayak, S. (2024). *Phonetic and acoustic analyses of sound-symbolic associations of a large pseudoword set across multiple domains of meaning.* Unpublished master's thesis, Pennsylvania State University, College of Medicine, Hershey, PA, USA.

Sidhu, D. M., Westbury, C., Hollis, G., & Pexman, P. M. (2021). Sound symbolism shapes the English language: The maluma/takete effect in English nouns. *Psychonomic Bulletin and Review*, *28*(4), 1390–1398. https://doi.org/10.3758/s13423-021-01883-3

Vaden, K., Halpin, H., & Hickok, G. (2009). *Irvine Phonotactic Online Dictionary* (p. available from http://www.iphod.com.).
